# The role of intraspinal sensory neurons in the control of quadrupedal locomotion

**DOI:** 10.1101/2021.12.21.473311

**Authors:** Katrin Gerstmann, Nina Jurčić, Severine Kunz, Nicolas Wanaverbecq, Niccolò Zampieri

## Abstract

From swimming to walking and flying, animals have evolved specific locomotor strategies to thrive in different habitats. All types of locomotion depend on integration of motor commands and sensory information to generate precise movements. Cerebrospinal fluid-contacting neurons (CSF-cN) constitute a vertebrate sensory system that monitors CSF composition and flow. In fish, CSF-cN modulate swimming activity in response to changes in pH and bending of the spinal cord, yet their role in higher vertebrates remains unknown. We used mouse genetics to study their function in quadrupedal locomotion and found that CSF-cN are directly integrated into spinal motor circuits by forming connections with motor neurons and premotor interneurons. Elimination of CSF-cN selectively perturbs the accuracy of foot placement required for skilled movements at the balance beam and horizontal ladder. These results identify an important role for mouse CSF-cN in adaptive motor control and indicate that this sensory system evolved a novel function from lower vertebrates to accommodate the biomechanical requirements of terrestrial locomotion.

## Introduction

Animals have developed a wide variety of locomotor strategies to adapt to their environment. The ability to precisely control movements is essential for each mode of locomotion and depends on the dynamic integration of motor commands and sensory information. Planning and initiation of motor programs take place in the brain, while their execution is directed by the spinal cord. Spinal circuits combine descending input with sensory feedback in order to generate coordinated movements and reflexive actions (Rossignol et al., 2006; Koch et al., 2018; Tuthill and Azim, 2018). While the contributions of the somatosensory system have been extensively studied, the role of other sources of sensory information is less clear.

CSF-cN have been first described a century ago as sensory neurons lining the central canal in vertebrates (Kolmer, 1921; Agduhr, 1922). They exhibit a peculiar morphology including a ciliated protrusion extending into the lumen of the central canal (Vigh et al., 2004). Thus, they have been thought to represent a sensory system monitoring CSF composition and flow. CSF-cN are inhibitory neurons and express ion channels known to be involved in sensory transduction, such as P2X and Pkd2l1 (Stoeckel et al., 2003; Orts-Del’Immagine et al., 2016). The latter represents a specific marker of CSF-cN (Djenoune et al., 2014). In larval zebrafish, CSF-cN are directly connected to primary motor neurons and V0v interneurons, glutamatergic premotor neurons that are part of the swimming central pattern generator (Fidelin et al., 2015, Hubbard et al., 2016). Optogenetic stimulation of CSF-cN in resting larvae elicits low frequency movements, while activation during active swimming results in inhibition of locomotion, thus indicating that CSF-cN differentially modulate motor activity depending on the state of the animal (Fidelin et al., 2015).

In fish, CSF-cN sense changes in pH and spinal curvature (Böhm et al., 2016; Jalalvand et al., 2016 a, Jalalvand et al., 2016 b). In particular, calcium imaging experiments revealed responses to both active and passive bending of the body axis, highlighting CSF-cN function as a mechanosensory system detecting the curvature of the spinal cord, either self-generated or induced by external forces (Böhm et al., 2016; Hubbard et al., 2016). Pkd2l1 has been shown to be crucial for CSF-cN mechanosensory function, in its absence CSF-cN are not activated by spinal bending and behavioral responses are impaired (Böhm et al., 2016; Sternberg et al., 2018). Altogether these studies indicate that CSF-cN are the key component of a chemo- and mechanoreceptive sensory system that relay information about CSF composition and curvature of the body axis in order to modulate locomotor activity and control posture (Orts-Del’Immagine and Wyart, 2017).

The biomechanical requirements and circuit mechanisms controlling wave-like propagation of swimming movements in fish versus on-ground locomotion in higher vertebrates are substantially different, raising questions regarding the physiological function of CSF-cN in terrestrial mammals (Grillner and Jessell, 2009). In this study, we analyzed CSF-cN connectivity and function in mice. We found that CSF-cN are connected to motor neurons and V0c interneurons, a key component of spinal motor circuits involved in the modulation of motor neuron firing frequency (Zagoraiou et al., 2009). CSF-cN ablation did not affect motor activity nor the generation of stereotyped locomotor patterns, such as walking and swimming, but resulted in specific defects in skilled locomotion. We observed an increase of foot slips and falls at the balance beam and the horizontal ladder, indicating that elimination of CSF-cN leads to a reduction in the accuracy of paw placement in unstable walking conditions. Surprisingly, we found that in mice Pkd2l1 activity is dispensable for CSF-cN function. However, elimination of CSF-cN cilium is sufficient to completely recapitulate the phenotypes observed after neuronal ablation, thus demonstrating that this structure is necessary for sensory transduction. Altogether this study shows that during the evolutionary transition from swimming to walking, CSF-cN have acquired a novel role in order to adapt to the specific needs of limbed-based locomotion, and are an essential part of to the sensory feedback mechanisms that control adaptive motor reflexes required for precise foot placement.

## Results

### CSF-cN connect to motor neurons and V0c premotor interneurons

We studied the physiological role of CSF-cN in the mammalian nervous system by obtaining selective genetic access using the *Pkd2l1::Cre* mouse line (Ye et al., 2015). We verified targeting specificity by lineage tracing with a nuclear GFP reporter line (*Rosa*Φ*HTB*; Li et al., 2013) and found that at all spinal levels ∼ 84% of labelled cells were Pkd2l1^+^ and presented the characteristic position and morphology of CSF-cN (Figures S1A and S1B). In addition, we did not observe reporter expression in any other cell type in the central nervous system (Figure S1C).

In order to explore CSF-cN connectivity, we first investigated synaptic targets by labelling pre-synaptic boutons with tdTomato-tagged synaptophysin (*Ai34*; Daigle et al., 2018). We observed dense signal localized around the central canal and in the ventromedial part of the spinal cord at all axial levels (Figure 1A). Interestingly, key components of spinal motor circuits are characterized by stereotyped positioning in these areas along the entire rostro-caudal extent of the spinal cord. Median motor column (MMC) neurons controlling the activity of epaxial muscles are found in ventromedial location and V0c neurons, cholinergic premotor interneurons, are positioned in the intermediate spinal cord on the sides of the central canal (Dasen and Jessell, 2009; Zagoraiou et al., 2009). Thus, we asked whether MMC and V0c neurons receive synaptic input from CSF-cN. To test this hypothesis, we relied on the cholinergic nature and stereotyped position of these cell types to identify them. We found putative synaptic contacts on ∼57% of V0c and ∼35% of MMC neurons at all spinal levels (Figures 1B, 1C and S1D). In addition, we also observed instances of tdTomato^+^ boutons juxtaposed to lateral motor column (LMC) neurons (∼13%; Figures 1C and S1D).

**Figure 1.**
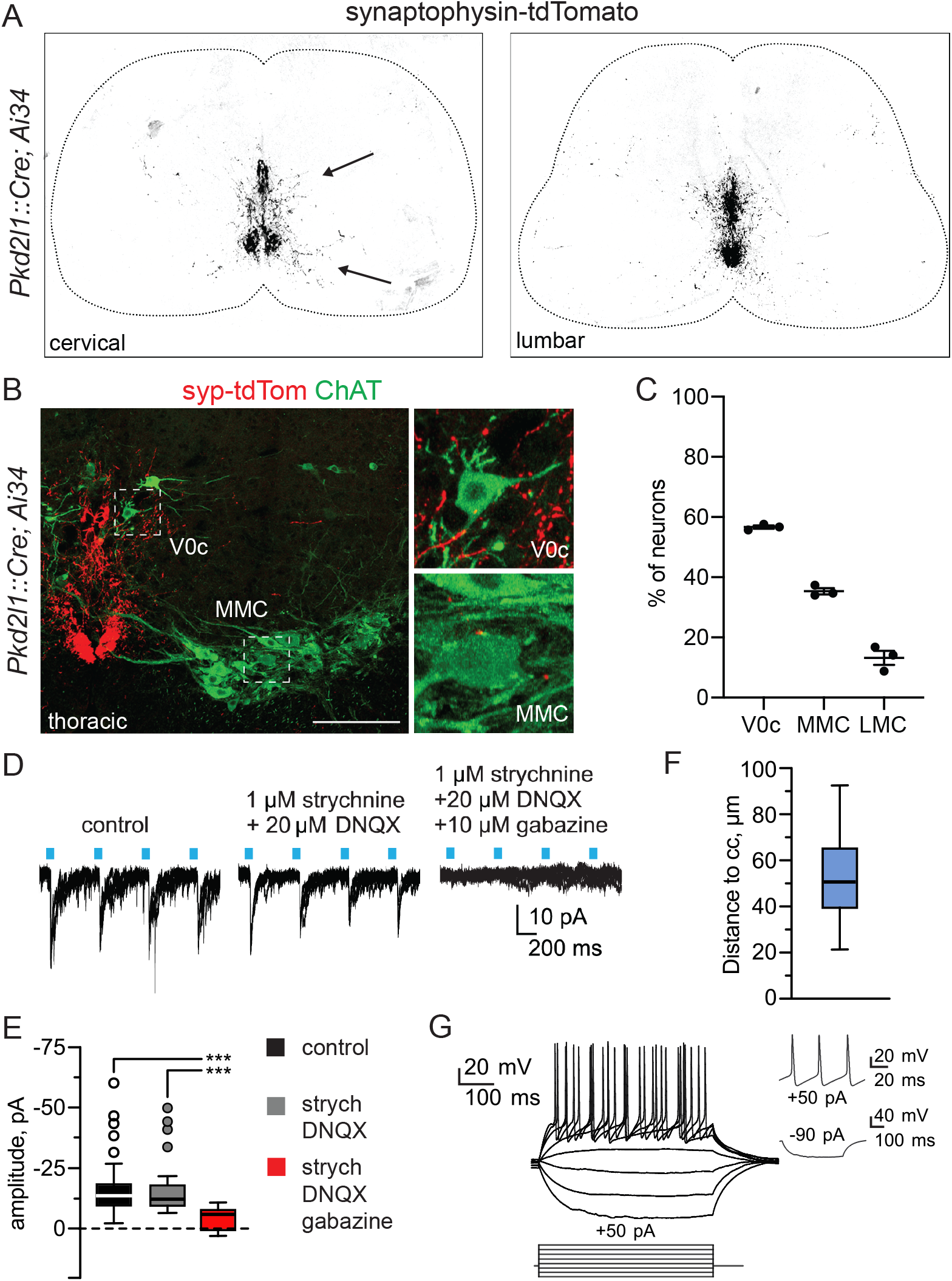
CSF-cN project to V0c and motor neurons. A) Representative images of synaptophysin-tdTomato labeling of CSF-cN at cervical and lumbar levels of P7 *Pkd2l1::Cre; Ai34* mice. Arrows point to dense labeling nearby the central canal and the ventromedial area of the spinal cord. B) Representative images of synaptophysin-tdTomato puncta in close contact with ChAT^+^ V0c and MMC neurons at thoracic level of P7 *Pkd2l1::Cre; Ai34* mice. C) Proportion of V0c, MMC and LMC neurons that receive synaptic input from CSF-cN (n=3). D) Optical activation of CSF-cN induce inhibitory currents in post-synaptic partners positioned nearby the central canal, that can be blocked by gabazine administration, but not with strychnine or DNQX. E) Amplitudes of inhibitory currents induced in post-synaptic partners after optical activation of CSF-cN. F) Distance to the central canal of responsive postsynaptic partners. G) Representative traces of electrical potentials in responsive post-synaptic partners after optical activation of CSF-cN. Mean±SEM, two-way ANOVA, ns p>0.05, *** p<0.001.

To confirm these anatomical findings and assess functional connectivity, we expressed channelrhodopsin-2 (ChR2, *Ai32*. Madisen et al., 2012) in CSF-cN and used whole-cell patch-clamp combined with ChR2-assisted circuit mapping to record from putative CSF-cN postsynaptic partners (Petreanu et al., 2007). In line with the inhibitory phenotype of CSF-cN, we found that short light pulses evoked inhibitory responses in neurons (20/500 neurons patched) located in the proximity of ChR2^+^ varicosities that were abolished in the presence of gabazine (Figures 1D and 1E). The majority of responsive neurons (15/20) were located in close proximity of the central canal (50 ± 5 µm) and presented electrophysiological (r_*m*_: 541 ± 7 MΩ; c_*m*_: 26 ± 2 pF) and action potentials properties (AP half width: 2.0 ± 0.2 ms; discharge frequency: 26 ± 2 Hz, +50 pA direct current injection) characteristic of V0c neurons (Figures 1F and 1G. Zagoraiou et al., 2009). The remaining cells were characterized by a different physiological profile suggesting that at least another interneuron subtype residing next to the central canal receive direct input from CSF-cN (Data not shown). Altogether, these data indicate that CSF-cN connect to motor neuron and V0c interneurons.

### CSF-cN are reciprocally connected and receive sparse input from spinal interneurons

Next, we explored sources of presynaptic input to CSF-cN by using rabies virus (RV) retrograde monosynaptic tracing (Wickersham et al., 2007). We selectively targeted CSF-cN for rabies infection and transsynaptic spread by injecting a mixture of Cre-dependent helper adeno-associated viruses at lumbar (L) level 1 of *Pkd2l1::Cre* mice (AAV-syn-FLEX-splitTVA-EGFP-tTA and AAV-TREtight-mTagBFP2-B19G; Lavin et al., 2020). Three weeks later, EnvA pseudotyped G-deficient RV (RVΔG-mCherry/EnvA) was delivered at the same level (Figure 2A). We first examined starter cells, defined as neurons infected by both AAV and RV, and found BFP^+^; RV^+^ neurons around the central canal at the point of injection, with morphologies and positions characteristic of CSF-cN (Figures 2B-D, S2A, S2C and S2D). We next focused on transynaptically labelled neurons and found that the majority of BFP^-^; RV^+^ cells were also CSF-cN (∼ 85%), but mainly located at more caudal levels of the spinal cord, thus indicating that CSF-cN are reciprocally connected, with caudal neurons sending input to rostral segments (Figures 2C, 2D and S2B-D). The remaining presynaptic input consisted of sparse labeling of spinal interneurons without any distinct positional organization (Figures 2C-E and S2B-D). Next, we analyzed the neurotransmitter phenotype of CSF-cN presynaptic partners by assessing expression of *VGAT* and *VGLUT2* to define inhibitory and excitatory status of second order neurons. We found that the majority of RV^+^ neurons were *VGAT*^+^ including CSF-cN that are known to have GABA-ergic phenotype (Figures 2F and S2E; Stoeckel et al., 2003). Altogether, these data indicates that CSF-cN are reciprocally connected and receive sparse input by spinal interneurons.

**Figure 2.**
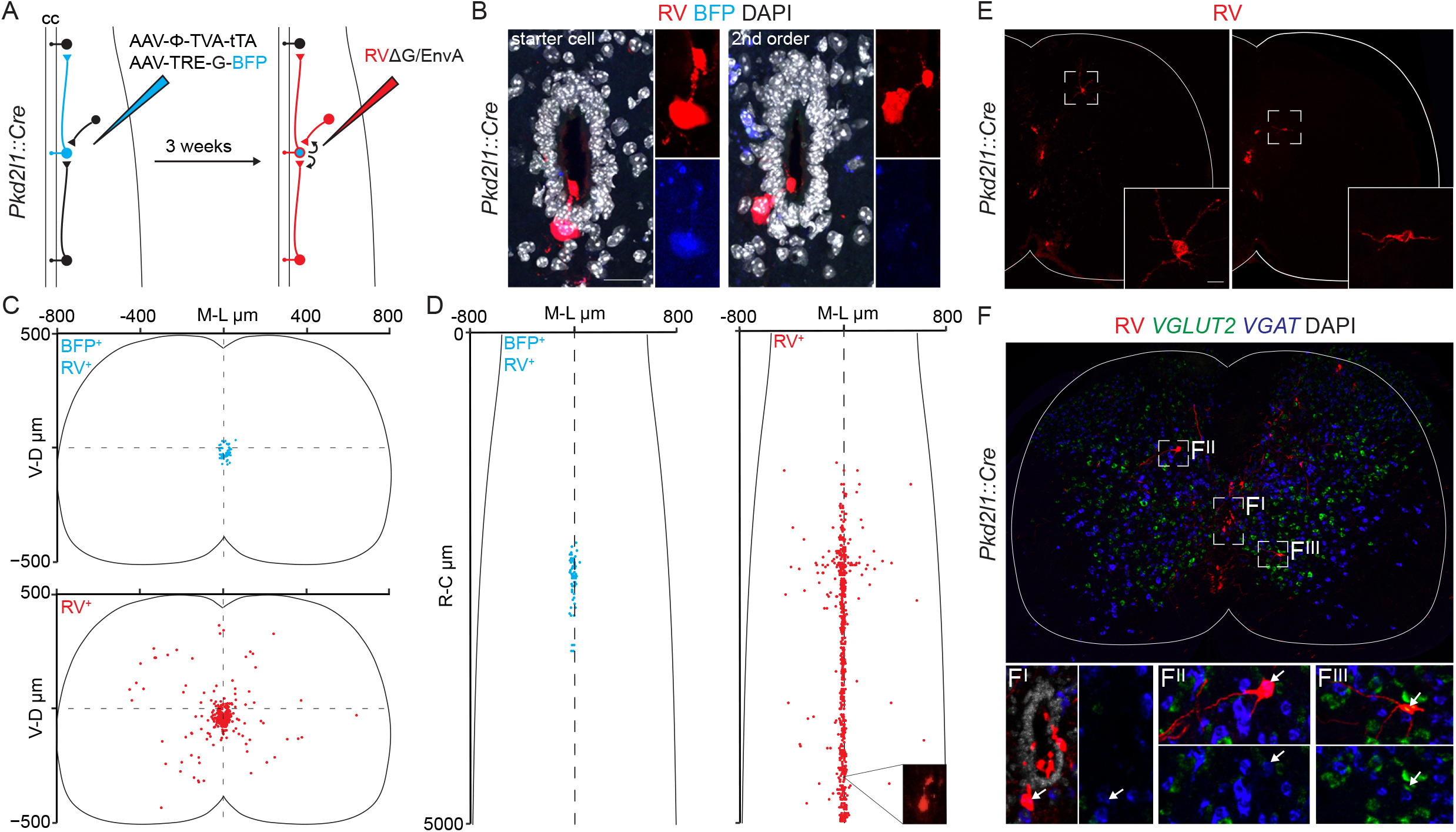
CSF-cN are reciprocally connected and receive sparse input from local spinal interneurons. A) Schematic illustrating rabies monosynaptic tracing approach to identify cells providing input to CSF-cN. A mix of Cre-dependent helper AAVs was injected at L1 of P7 *Pkd2l1::Cre* mice. Three weeks later, RVΔG-mCherry/EnvA was injected at the same position and after 7 days spinal cords examined. B) Representative images of BFP^+^; RV^+^ starter cells and BFP^-^; RV^+^ second order CSF-cN. Scale bar: 20 µm. C-D) Digital reconstruction of medio-lateral/dorso-ventral position (C) and medio- lateral/rostro-caudal position (D) of starter cells (blue) and second order cells (red); n=3. E) Representative images of second order neurons labeled in rabies tracing experiments from CSF-cN. F) Multiplexed fluorescent in situ hybridization analysis of RV^+^ neurons showing excitatory (VGLUT2^+^) and inhibitory (VGAT^+^) second order cells.

### CSF-cN are required for accurate foot placement during skilled walking

The organization of mouse CSF-cN input and output connectivity suggests a role in motor control. To study the function of CSF-cN, we crossed the *Pkd2l1::Cre* line with the *Rosa*Φ*DTR* allele to drive expression of the diphtheria toxin receptor (DTR; Buch et al., 2005). Diphtheria toxin (DT) administration in adult mice resulted in elimination of >80% CSF-cN within two weeks (Figures 3A, 3B, S3A and S3B). We first evaluated the effect of acute ablation of CSF-cN on general locomotor function associated with exploratory behavior by assessing performance in the open field test. We did not find any significant difference between DT- and PBS-treated mice in activity, speed, or distance travelled (Figures 3C-E). Next, we performed kinematic analysis on freely walking mice to evaluate gait and did not observe any effect on step cycle, step length, and base of support, key parameters describing limb movement and coordination (Figures 3F, 3G, S3C and Video S1; Mendes et al., 2015). These data show that elimination of CSF-cN does not perturb locomotor activity and the generation of the patterns and rhythms of muscle contraction necessary for walking gait in mice. In larval zebrafish, CSF-cN have been shown to have an important role for postural control (Hubbard et al., 2016). Thus, we challenged the mice with tasks requiring precise control of trunk position and stability. First, we evaluated spontaneous rearing events, and found no effect of CSF-cN ablation on rearing duration and frequency (Figure S3D and data not shown). Second, we tested swimming, a locomotor behavior that requires coordination of limb and trunk muscles in order to obtain directional movements (Gruner and Altman, 1980). We did not observe any difference in speed or in the angle between the trunk and the water line, an indicator of postural control (Figures S3E, S3F and Video S2; Pocratsky et al., 2020). These experiments show that elimination of CSF-cN does not perturb postural control.

**Figure 3.**
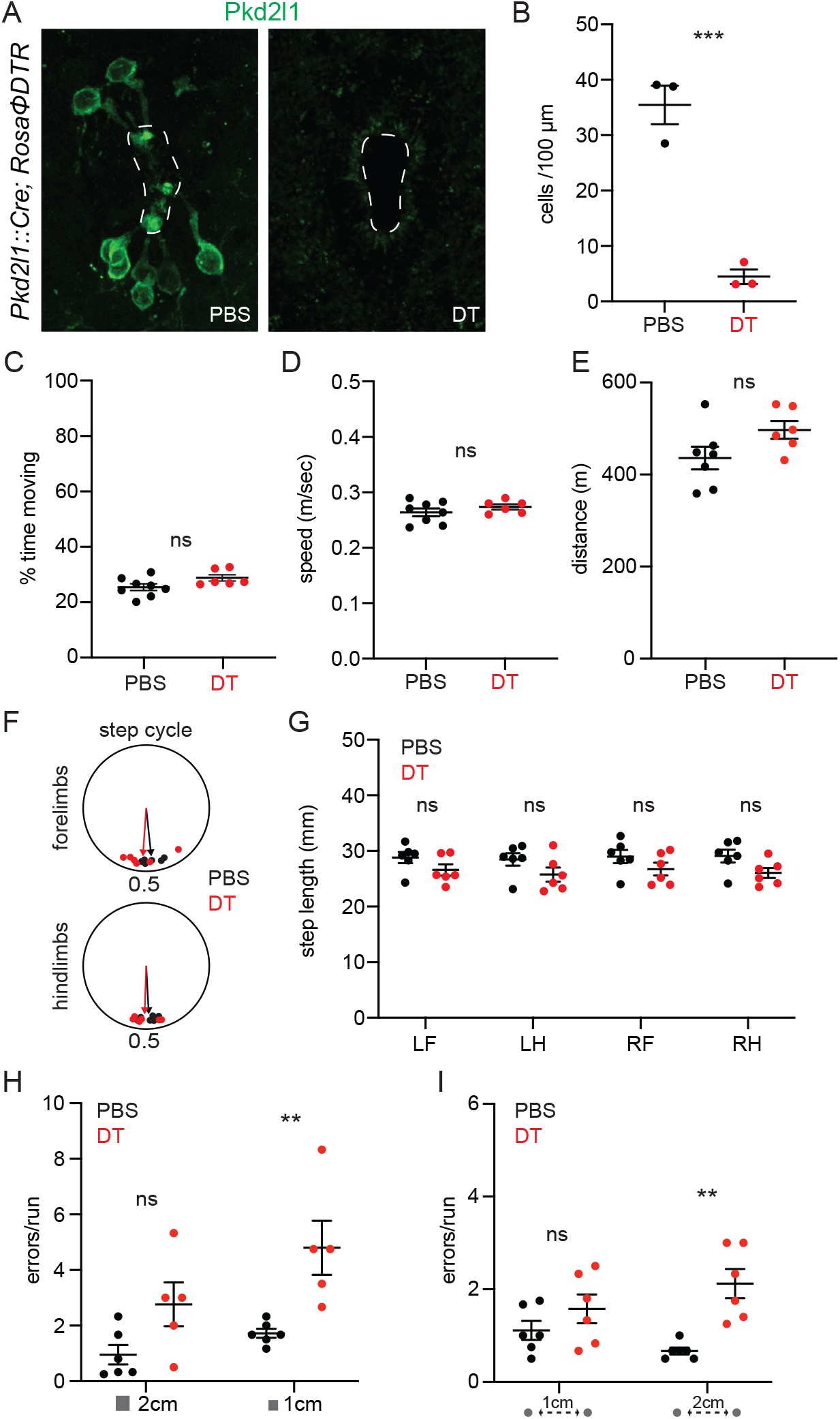
Pharmacological ablation of CSF-cN perturbs skilled locomotion. A) Representative images of Pkd2l1^+^ neurons around the central canal 60 days after PBS (left) or DT (right) injection in *Pkd2l1::Cre; Rosa*Φ*DTR* mice. B) Quantification of Pkdl21^+^ neurons around the central canal 60 days after PBS (n=3) or DT (n=3) injection in *Pkd2l1::Cre; Rosa*Φ*DTR* mice. C-E) Locomotor activity during a 90 min open field test. Percentage of time spent moving (C), speed (D) and distance traveled (E) in adult *Pkd2l1::Cre; Rosa*Φ*DTR* mice 14 days after PBS (n=8) or DT (n=6) treatment. F) Step cycle in adult *Pkd2l1::Cre; Rosa*Φ*DTR* mice 14 days after PBS (n=6) or DT (n=6) treatment. G) Step length in adult *Pkd2l1::Cre; Rosa*Φ*DTR* mice 14 days after PBS (n=6) or DT (n=6) treatment (LF left forelimb, LH left hindlimb, RF right forelimb, RH right hindlimb). H) Quantification of foot placement errors (slips and falls) in the balance beam test with 2 cm (left) or 1 cm (right) beam width in adult *Pkd2l1::Cre; Rosa*Φ*DTR* mice 14 days after PBS (n=6) or DT-injection (n=5). I) Quantifications of foot placement errors (slips and falls) in the horizontal ladder test with 1 cm (left) or 2 cm (right) rung distance with adult *Pkd2l1::Cre; Rosa*Φ*DTR* mice 14 days after PBS (n=6) or DT-injection (n=6). Mean±SEM, paired t-test, ns p>0.05, ** p<0.01, *** p<0.001.

Finally, we tested skilled locomotion by assessing walking on the balance beam and horizontal ladder, tasks that are known to require tactile and proprioceptive sensory feedback in order to achieve regularity and accuracy in foot placement (Akay and Murray, 2021). We used beams and ladders of different widths (2 cm and 1 cm) and rung spacing (1 cm and 2 cm) in order to assess the effect of progressively more difficult conditions (Rossignol et al., 2006). In both tasks, DT-treated mice presented clear deficits in motor performance (Videos S3 and S4) that resulted in increase in the numbers of foot slips and falls, which was significantly higher than control animals in the more challenging configurations (Figures 3H and 3I). Thus, these data indicate that CSF-cN contribute to the control of adaptive movements required for precise foot placement during skilled locomotion.

### The cilium is necessary for CSF-cN function

Next, we wondered whether the Pkd2l1 channel is necessary for CSF-cN function in mice, as in zebrafish its elimination impairs mechanosensation and behavioral responses to changes in spinal bending (Böhm et al., 2016; Sternberg et al., 2018). To address this question, we analyzed locomotor behavior in Pkd2l1 knockout mice (*Pkd2l1-/-*; Figure S3G. Horio et al., 2011). In line with the results obtained after neuronal ablation experiments these mice did not show any phenotype at the open field, gait analysis, and swimming tests (Figures S3H and S3I; Videos S1 and S2). Surprisingly, *Pkd2l1-/-* mice performance at the balance beam and horizontal ladder was also indistinguishable from control mice (Figures S3J and S3K; Videos S3 and S4). Altogether these data indicate that CSF-cN role in the control of adaptive motor reflexes in mammals does not require Pkd2l1 activity.

Cilia have been known to function as a mechanosensory organelle responding to fluid flow in many different cell types and most notably in sensory neurons. Thus, we studied the consequences of preventing cilium formation in CSF-cN. The intraflagellar transporter 88 (Ift88) is part of the Ift-B complex that is crucial for transport of ciliary proteins and its elimination suppress ciliogenesis (Pazour et al., 2000). We prevented cilium formation in CSF-cN by crossing the *Pkd2l1::Cre* allele with the conditional *Ift88*^*fl*^ mouse line (*Pkd2l1::Cre +/-; Ift88*^*fl/fl*^, hereafter referred to as Δ*Cilia*. Haycraft et al., 2007). We first confirmed success of this strategy by visualizing CSF-cN protrusions in the central canal and the associated cilium (Figure 4A). In control animals, we found that >70% of Pkd2l1^+^ apical processes presented a cilium, while in Δ*Cilia* mice we found a significant reduction in the occurrence of ciliated CSF-cN (∼35%; Figures 4A and 4B). Moreover, electron microscopy analysis confirmed that conditional elimination of Ift88 prevents ciliogenesis in CSF-cN (Figures S4A and S4B). Next, we evaluated locomotor behavior in Δ*Cilia* mice. We did not observe any significant defect in the open field, gait analysis, and swimming tests (Figures 4C-E and S4C-G; Videos S1 and S2). However, we observed that the performance of Δ*Cilia* mice at the balance beam and horizontal ladder was reminiscent of the one of mice lacking CSF-cN (Videos S3 and S4). Strikingly, quantification of foot slips and foot falls revealed that Δ*Cilia* made significantly more mistakes when walking on the more challenging versions of the tests, thus precisely recapitulating the phenotype observed after CSF-cN ablation (Figures 4F and 4G). Altogether these data show that elimination of cilium phenocopies neuronal ablation, thus indicating that the cilium, but not Pkd2l1 activity, is necessary for CSF-cN role in adaptive motor control in mice.

**Figure 4.**
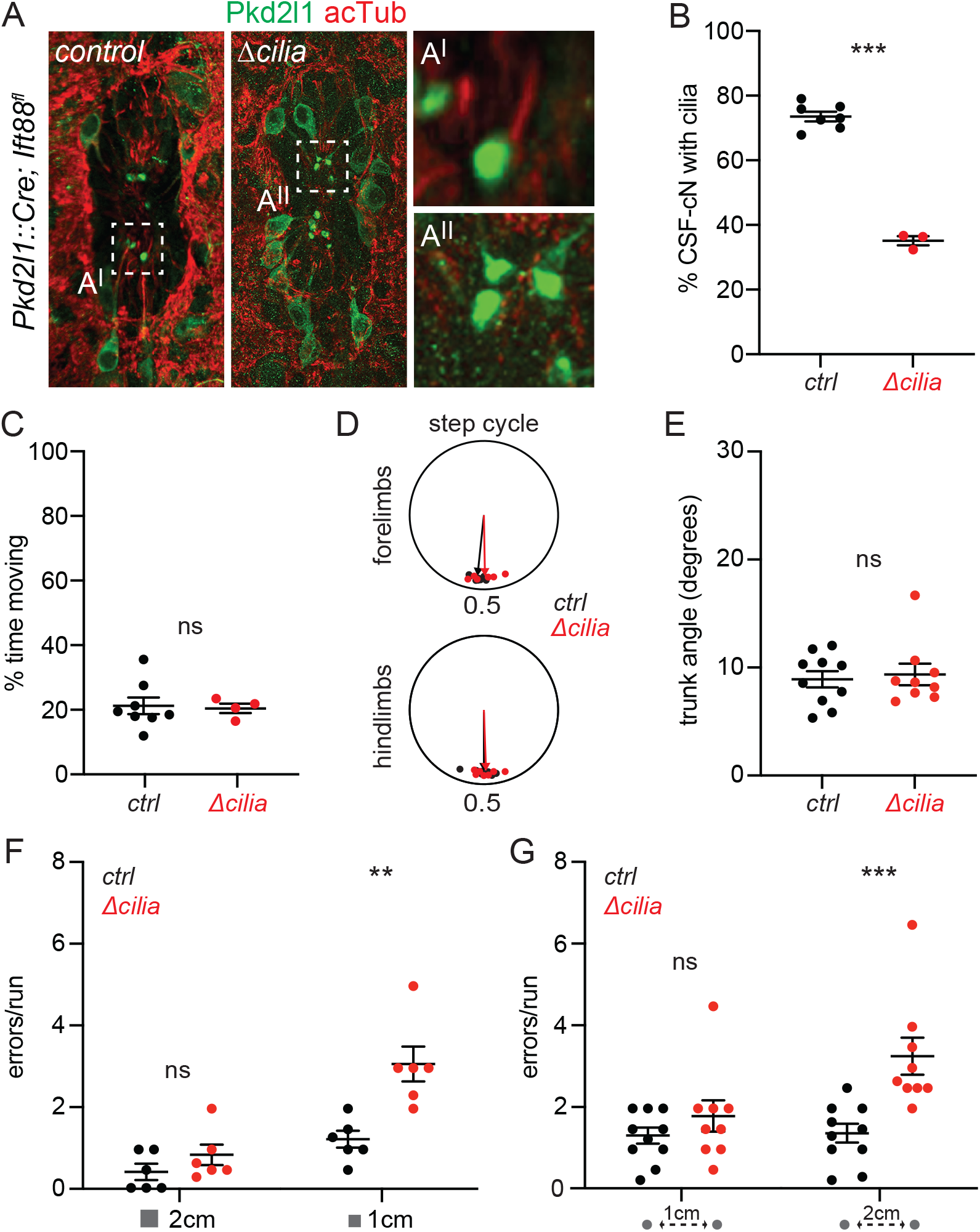
Elimination of CSF-cN cilium phenocopies neuronal ablation. A) Representative images of Pkd2l1^+^ CSF-cN apical protrusions and acetylated-Tubulin^+^ cilia in adult *control* and Δ*cilia* mice. High magnifications of Pkd2l1^+^ protrusion in the central canal of *control* (A^I^) and Δ*cilia* (A^II^) mice. B) Quantification of Pkd2l1^+^ CSF-cN apical protrusions bearing an acetylated-Tubulin^+^ cilium in *control* and Δ*cilia* mice (n=3). C) Percentage of moving time in adult *control* (n=7) and Δ*cilia* mice (n=4) during 90 min open field test. D) Step cycle of forelimbs (top) and hindlimbs (bottom) in adult *control* (n=10) and Δ*cilia* mice (n=9). E) Quantification of trunk angle between body axis and water line during swimming task in adult *control* (n=10) and Δ*cilia* mice (n=9). F) Quantification of foot placement errors (slips and falls) in the balance beam test with 2 cm (left) or 1 cm (right) beam width in adult *control* (n=6) and Δ*cilia* mice (n=6). G) Quantification of foot placement errors (slips and falls) in the horizontal ladder test with 1 cm (left) or 2 cm (right) rung distance in adult *control* (n=10) and Δ*cilia* mice (n=9). Mean±SEM; paired t-test, ns p>0.05, ** p<0.01, *** p < 0.001.

## Discussion

In this study, we investigated the physiological role of CSF-cN, an evolutionary conserved vertebrate sensory system, in limbed mammals. We found that these neurons are integrated into spinal motor circuits and contribute to adaptive motor reflexes at the basis of accurate foot placement during skilled locomotion. CSF-cN function does not require the activity of the Pkd2l1 channel but entirely depends on its cilium, thus pointing to a key role for this mechanosensory structure in monitoring CSF flow. Altogether, our data suggests a model where CSF-cN provide an additional source of proprioceptive information by monitoring spinal curvature and represent an important component of the sensory feedback mechanisms ensuring accuracy in motor control.

Kolmer and Agduhr first described a peculiar population of sensory neurons lining the central canal and proposed that they constitute a sensory organ relaying information from the CSF (Kolmer, 1921; Agduhr, 1922). CSF-cN function has remained elusive until recent studies in lower vertebrates revealed physiological roles in modulation of locomotion and postural control (Fidelin et al., 2014; Jalalvand et al., 2016; Böhm et al., 2016). CSF-cN have been shown to provide information to the motor system about active and passive curvature of the body axis by sensing fluid flow along the central canal. (Böhm et al., 2016; Sternberg et al., 2018; Orts-Del Immagine et al., 2020). In fish, the importance of monitoring curvature along the rostro-caudal axis of the spinal cord is clear, as swimming relies on the rhythmic propagation of an undulatory pattern of muscle contraction. The introduction of limbs in higher vertebrates have led to the reorganization of motor circuits in order to accommodate the biomechanical requirements of terrestrial locomotion (Grillner and Jessell, 2009). The regularity and precision of foot placement represent a critical feature of motor control in limbed animals (Grillner and El Manira, 2020). In particular, it is especially important for navigating the diverse terrains and obstacles animals are confronted with in the wild and require dynamic integration of different sources of sensory information (Akay and Murray, 2021). Thus, our data suggests that the role of CSF-cN as a sensory system monitoring curvature of the body axis has evolved from modulation of wave-like tail movements at the basis of swimming in fish, to the control of adaptive motor responses necessary to regulate the accuracy of foot placement in limbed mammals.

At circuit level, we observed connectivity patterns that are consistent with CSF-cN physiological role in sensorimotor integration. We provide evidence that CSF-cN have direct and indirect access to spinal motor circuits. First, we found CSF-cN presynaptic puncta on MMC neurons, which selectively innervate epaxial muscles, thus providing a direct way to regulate trunk movements. Sparse innervation on limb innervating motor neurons was also observed, offering the ability to directly influence limb movements. Second, dense CSF-cN input was found on V0c premotor interneurons, thus in principle providing indirect access to the activity of multiple muscle groups at once, as single V0c have been shown to innervate functionally equivalent motor neurons on both sides of the spinal cord (Stepien et al., 2010). V0c neurons are known to modulate the intensity of motor output in a task-dependent manner and are therefore in prime position to tune the firing of individual muscles in order to adapt to the requirements of skilled locomotion (Zagoraiou et al., 2009; Witts et al., 2014). However, how V0c activity is adjusted to match the needs of different locomotor activities is not known. Sensory feedback has been proposed to be a possible mechanism and indeed V0c receive input from primary somatosensory neurons and dorsal spinal interneurons, thus representing an ideal node for integrating different sources of sensory information in order to regulate muscle activity in a context-dependent manner (Zampieri et al., 2014).

In addition, we provide evidence that CSF-cN receive sparse input from interneurons located at the same segmental level, mostly of inhibitory character. Inhibitory neurons could control the gain of CSF-cN activity, a well-known mechanism for tuning somatosensory feedback in spinal circuits (Rudomin and Schmidt, 1999). Finally, we observed CSF-cN recurrent connectivity in the rostral direction, zebrafish CSF-cN are known to send ascending projections up to targets in the midbrain (Wu et al., 2021). An inhibitory feedback loop from posterior CSF-cN to anterior ones is well suited for coordination of undulatory movements in fish, but its significance for limb-based locomotion remains to be explored but could be exploited for coordinating muscle activity at different segmental levels.

At a functional level, our data show that in mice elimination of CSF-cN does not perturb general activity and the generation of rhythmic patterns of limb movement necessary for the production of stereotyped locomotor actions, such as walking and swimming, but have a specific role in the control of corrective motor reflexes. In absence of CSF-cN, we observe an increase of foot slips and falls at the balance beam and horizontal ladder, indicating that the precision of paw placement required for performing skilled movements is perturbed. Interestingly, the effect is significant only in the most challenging versions of these tasks. These data highlights how multiple sources of sensory information, including cutaneous and muscle afferents, the visual, and the vestibular systems, are integrated to control the postural adjustments and adaptive limb movements that are necessary to prevent foot slippage during the execution of skilled actions (Rossignol et al., 2006).

Previous work in lamprey and zebrafish, along with our observation that elimination of the cilium completely recapitulates the defects observed after neuronal ablation point to a main role for CSF-cN in mice as mechanoreceptive sensory neurons detecting bending of the spinal cord by sensing CSF flow in the central canal (Orts-Del’Immagine et al., 2020). Thus, we propose that CSF-cN by monitoring spinal bending provide proprioceptive feedback informing the motor system on axial position that is used to adjust trunk movement and paw placement during locomotion. Walking on narrow paths or challenging terrains introduces forward and lateral displacements in the body axis that can be finely monitored by CSF-cN. For example, walking on a balance beam reduces lateral stability by decreasing the available base of support or walking on the horizontal ladder requires to overextend hindlimbs in order to land on the same rung where the forelimbs touched down, thus resulting in exaggerated hip torsion and spinal bending (Akay and Murray, 2021).

Surprisingly we did not observe any defect in locomotor behavior upon elimination of Pkd2l1. This is in contrast with its requirement for CSF-cN function in zebrafish, thus raising interesting questions regarding molecular effectors in mammals. Our study does not exclude the possibility that in mice Pkd2l1 might be selectively required for chemosensation, as this channel has been shown to respond directly to pH changes (Orts-Del’Immagine et al., 2015). The ability to monitor CSF composition has been proposed to be part of a homeostatic mechanisms common to all vertebrates for counteracting the effects of pH changes by reducing muscle activity (Jalalvand et al., 2016 a; Jalalvand et al., 2016 b). It will be interesting to address whether chemosensation in mammalian CSF-cN could serve as system for modulating motor behavior in response to changes in the internal state of the animal, for example in case of fatigue or sickness.

Altogether, our anatomical and functional data indicate that CSF-cN are an important component of sensorimotor circuits in the mammalian spinal cord contributing to adaptive motor control. This study opens the way for future work to address exciting questions on how information on CSF composition and flow is encoded by CSF-cN and integrated at a circuit level with other sensory input, such as muscle and cutaneous feedback, in order to orchestrate flawless execution of motor programs.

## Acknowledgements

We thank Liana Kosizki for technical support and the MDC Advanced Light Microscope facility for assistance with image acquisition and analysis. We thank Robert Manteufel, Ilka Duckert and Florian Keim for animal care; Baptiste Lasbats and Lilly von Kalckreuth for assistance with behavioral experiments; Pierre-Louis Ruffault for advice with experimental design; Nikos Balaskas, Marco Beato, Joriene De Nooij and members of the Zampieri laboratory for insightful comments on the manuscript. N.Z. and NW were supported by a DFG-ANR international collaborative grant (MotAct-CSF. DFG ZA 885/1-2; ANR-16-CE92-0043); N.W. by AMU and CNRS INSB.

## Author contributions

Conceptualization, K.G., N.W., and N.Z.; Investigation, K.G., N.J., and S.K.; Formal analysis, K.G., N.J., and S.K.; Writing – Original Draft, K.G. and N.Z.; Writing – Review and Editing, K.G., N.W., and N.Z.; Supervision, N.W., and N.Z.

## Material and Methods

### Animal Experimentation Ethical Approval

All animal procedures were performed in accordance to European Communities Council Directives and were approved by the Regional Office for Health and Social Affaires Berlin (LAGeSo) under license number G148/17.

### Animal models

Mice were bred and maintained under standard conditions on a 12h light/dark cycle with access to food and water *ad libitum*. The day of birth was considered as postnatal day 1 (P1).

### Ablation of CSF-cNs

To specifically ablate CSF-cNs *in vivo*, diphtheria toxin (DT; Sigma D0564) was administered intraperitoneally (50mg/kg) at P40. Ablation efficiency was verified by staining for Pkd2l1.

### Behavioral experiments

Mice were placed in the behavior room 30 min before starting the experiments, allowing them to acclimatize. Both sexes were included and for each test at least two representative videos with continuous movements were analyzed. For the *open field* test we used the ActiMot Infrared light beam activity monitor (TSE Systems). Two light-beam frames allowed to monitor X, Y and Z coordinates of the mouse. Animals were placed in the associated squared acrylic glass boxes (40 cm X 20 cm) and after 10 min of habituation time, spontaneous movements were monitored for 90 min. Data were evaluated with TSE supplied software. *Gait analysis* was performed as previously described (Mendes et al., 2015). Briefly, mice were placed on a customized acrylic glass walkway with surrounded LED lights to generate the internal reflection effect. A mirror under the walkway allows tracking of footprints and body outline with a high-speed camera (shutter speed 5,56 ms, frame rate 150 f/sec). Representative videos with straight and continuous runs were analyzed using the open-source MouseWalker software. To evaluate balance, we used a customized *balance beam* with replaceable beams of different sizes. Animals were placed on one end and had to pass the beam spontaneously to reach a shelter on the other side. A mirror was placed underneath and a high-speed camera captured the passage. The *horizontal ladder* was customized with side walls made of acrylic glass to create a walking path and inserted metal rungs with 3 mm diameter. Rungs had a minimum distance of 1 cm and spacing of the rungs were modified by removing individual rungs. A mirror under the horizontal ladder and the clear walls allowed tracking from the side and underneath with a high-speed camera. Animals were required to pass the walking floor spontaneously and videos with continuous runs were analyzed. For the *swim task*, a custom-build acrylic glass tank (10 cm X 70 cm) filled with ambient temperature water was used. Mice had to swim through the tank to reach a platform on the other end. A mirror underneath allowed monitoring swim movements with a high-speed camera. The angle between body axis and water line was obtained by using the open-source program DeepLabCut (Mathis et al., 2018). The algorithm was trained to extract coordinates of nose and tail base in all frames. A value of likelihood allowed to estimate the reliability of detected coordinates and only frames with a likelihood superior to 0.9 were used for further analysis. The x/z coordinates of indicated points allowed the calculation of the swim angle between waterline and body axis.

### Perfusion and tissue preparation

Anesthesia was induced by the intraperitoneal injection of ketamine (120 mg/kg) and Xylazine (10 mg/kg). After testing the toe-pinch reflex, animals were intracardially perfused with 10 ml ice-cold PBS, followed by the perfusion of ice-cold 4 % PFA (pH 7.4). The spinal cords were exposed via laminectomy and post-fixed overnight in 4 % PFA (pH 7.4) at 4 °C. After washing for 5 min in PBS, tissue was incubated in 30% sucrose over night at 4 °C for cryoprotection. Samples were embedded in Optimal Cutting Temperature (O.C.T., Tissue-Tek) compound, frozen on dry ice and stored at -80 °C.

### Slice preparation, electrophysiology and optogenetic

*Pkd2l1::Cre; Rosa*Φ*ChR2*(Ai32) mice (3 to 4 weeks old) were deeply anesthetized with an intraperitoneal injection of a Ketamine/xylazine mixture (120/10 mg/kg) and perfused intracardiacally with an ice cold and oxygenated (95% O_2_/5% CO_2_) modified artificial cerebrospinal fluid (aCSF, in mM: NaCl 75, NaH_2_PO_4_ 1.25, NaHCO_3_ 33, KCl 3, MgSO_4_ 7, sucrose 58, glucose 15, ascorbic acid 2, myo-inositol 3, sodium pyruvate 2, CaCl_2_ 0.5, pH 7.4, 310 mosmol.kg-1). Following laminectomy and spinal cord extraction, lumbar spinal cord coronal slices (250 µm) were prepared, transferred in a submerged incubation chamber filled with oxygenated aCSF (in mM: NaCl 115, NaH_2_PO_4_ 1.25, NaHCO_3_ 26, KCl 3, MgSO_4_ 2, glucose 15, ascorbic acid 2, myo-inositol 3, sodium pyruvate 2, CaCl_2_ 2; pH 7.4, 300 mosmol.kg-1) at 35° C for 15 min and then for 1 h at room temperature (20-25°C). Slices were transferred in the recording chamber perfused at 2-4 mL/min with aCSF at room temperature (20-22°C). Electrodes (3-6 MΩ, borosilicate glass Harvard Apparatus) were filled with a solution containing (in mM): K-gluconate 120, NaCl 5, HEPES 10, MgCl_2_ 1, CaCl_2_ 0.25, EGTA 2, Mg-ATP 4, Na_2_-phosphocreatine 10, Na_3_-GTP 0.2 (pH 7.3, 295 mosmol.kg^-1^) and 20 µM AlexaFluor488 (Invitrogen). Whole-cell patch-clamp recordings from identified interneurons 5IR_DIC) were made in voltage- and current-clamp mode using a MultiClamp 700B amplifier (Molecular Device Inc.). Synaptic currents were photo-elicited with short light pulses delivered through the objective (60x, LED: 490 nm, 2.2 mW.mm^-2^ for 5-10 ms) in control and in the presence of 1 µM strychnine (Sigma-Aldrich), 20 µM DNQX and 10 µM gabazine (BioTechne, UK).

### Immunohistochemistry

For histology, spinal cords were sectioned with a cryostat (Leica) and collected on Superfrost Plus® microscope slides (Thermo Fisher Scientific). Primary and secondary antibodies were diluted in 4 % BSA in 0.3 % TritonX in PBS. Slides were mounted with Vectashield (Vector). The following primary antibodies dilutions have been used: rabbit anti-Pkd21l1 (1/200; Millipore), goat anti-ChAT (1/200; Millipore), rabbit anti-dsred (1/1000; TaKaRa), mouse anti-acetylated Tubulin (1/500, Sigma), and chicken anti-GFP (1/1000; abcam). Images were taken with a confocal laser scanning microscope LSM800 (Zeiss).

### Surgical procedures

*Intraspinal injections* were performed as previously described (Zampieri et al., 2014). For analgesia, mice were subcutaneously injected with 5 mg/kg Carprofen 30 min before surgery. Anesthesia was induced with continuous inhalation of isoflurane (2-3 %) in oxygen (1.5 %), using an isoflurane vaporizer (Parkland Scientific). Dorsal laminectomy was performed to expose the lumbar spinal cord prior to virus injection using a pulled borosilicate glass pipette (World Precision Instruments, Inc.) and a micro syringe pump injector (Smart Touch). A total amount of 300 nl of virus was inoculated into two adjacent spots bilaterally 40 µm left and right to the midline. For AAV, a cocktail of AAV-TREthight-mTag BFP2-B19G (4.48*10^12^ VG/mL) and AAV-FLEX-SPLIT TVA-EGFP-tTA (5.79*10^10^ VG/mL) was injected. After three weeks we performed spinal injection of 300 nl RVΔG(EnvA)-mCherry at the same position and animals were sacrificed seven days after. Mice which postmortem revealed low viral labeling or a spread into the central canal were excluded from analysis.

### Fluorescent in situ hybridization

For mRNA detection via multiplex RNAscope, a modified protocol from Advanced Cell Diagnostics (ACD, 322360-USM) was used. Briefly, fixed spinal cord tissue was prepared and sectioned as described before. Spinal sections were post-fixed in 4 % PFA (pH 7.4) at 4 °C for 15 min. After washing and dehydration (at 4 °C in 50%, 70% and 100% Ethanol), a hydrophobic barrier was created around sections. After incubation with 3 % hydrogen peroxide solution (H_2_O_2_) at RT for 15 min, Protease IV treatment followed for 30 min at RT. C2 and C3 probes were dilutes 1/50 in sample diluent and hybridized for 2 hours at 40°C in a humid chamber in a HybEZ oven. For signal amplification and detection the RNAscope 2.5 HD Reagents Detection Kit-RED (ACD, 32360) was used according to the manufacturer’s instructions. After Detection of each channel, immunostaining was performed as described before and slices were mounted with ProLong Gold.

### Positional analysis

Three-dimensional positional analysis was performed as previously described (Dewitz et al., 2018). Spinal cords were sectioned in 40 µm slices and cartesian coordinates of spinal neurons per section were obtained using the imaging software IMARIS. Data were normalized to account for differences in spinal cord size and shape. The position of each neuron was digitally reconstructed by plotting the data in ‘R’ (R Foundation for Statistical Computing, Vienna, Austria, 2005), using a customized script. Correlation analysis have been done using the ‘‘corrplot’’ package to calculates the comparability of experiments using the Pearson correlation coefficient. Datasets were clustered hierarchically.

### Electron microscopy

Mice were perfused with 4 % (w/v) paraformaldehyde in 0.1 M phosphate buffer. Spinal cord was dissected and 2-3 mm^3^ cubes were fixed by immersion in 4 % (w/v) paraformaldehyde and 2.5% (v/v) glutaraldehyde in 0.1 M phosphate buffer for 2 hours at room temperature (RT). Samples were postfixed with 1% (v/v) osmium tetroxide for 3 hours at RT, dehydrated in a graded series of ethanol, and embedded in PolyBed® 812 resin (Polysciences, Germany). Ultrathin sections (60-80 nm) were stained with uranyl acetate and lead citrate, and examined at 80 kV with a Zeiss EM 910 electron microscope (Zeiss, Germany). Acquisition was done with a Quemesa CCD camera using iTEM software (Emsis GmbH, Germany).

### Statistical analysis

For behavior experiments, mice were randomly allocated into different experimental groups and data have been randomized before analysis whenever it was possible. Quantifications represent the average of at least three biological replicates per condition. Each dot represents one animal and error bars in all figures represent mean ± SEM. Number of samples (n) and the applied statistical test used for individual experiments are indicated in the figure legends. Significance was defined as * p<0.05; ** p<0.01; *** p<0.001. Statistical analyses were performed using Microsoft Excel or GraphPad Prism (Version 8, GraphPad Software).

## Supplemental Information Titles and Legend

***Figure S1. Specificity of CSF-cN genetic targeting***.

A) Representative image of Pkd2l1^+^; nuclear GFP^+^ neurons around the central canal of *Pkd2l1::Cre; Rosa*Φ*HTB* mice.

B) Quantification of Pkd2l1^+^; nuclear GFP^+^ neurons at cervical, thoracic, and lumbar spinal segments (cervical 182/220 cells, thoracic 198/230 cells, lumbar 179/209 cells, n=4; Mean±SEM).

C) Tomato expression in the brain of P7 *Pkd2l1::Cre; Rosa*Φ*tdTomato* mice (cortex, Ctx; ventricular zone, VZ; cerebellum, CB). Scale bar: 1mm.

D) Representative pictures of synaptophysin-tdTomato synapses on ChAT^+^ V0c, MMC, and LMC neurons at cervical and lumbar segments of P7 *Pkd2l1::Cre; Ai34* mice.

***Figure S2. Rabies monosynaptic tracing from CSF-cN***.

A) Digital reconstruction of BFP^+^; RV^+^ starter cells (blue) position. Each plot represents one animal.

B) Digital reconstruction of second order cells (red) position. Each plot represents one animal.

C) Cross correlation analysis of starter cells (left) and second order cells (right) positional coordinates. The scale indicates correlation values.

D) Number of starter cells (blue), second order cells (red) and second order CSF-cN in individual rabies tracing experiments (right, black).

E) Proportion of VGLUT2^+^ and VGAT^+^ second order neurons (excluding CSF-cN).

Mean ± SEM, paired t-test, * p<0.05.

***Figure S3. Behavioral analysis of CSF-cN ablated and Pkd2l1 mutant mice***.

A) Representative images of Pkd2l1 staining in adult *Pkd2l1::Cre; Rosa*Φ*DTR* mice 7 days after PBS (left) or DT-injection (right).

B) Percentage of Pkd2l1^+^ cells in respect to the PBS control at 7 days, 11 days and 60 days after DT treatment in adult *Pkd2l1::Cre; Rosa*Φ*DTR* mice.

C) Base of support of forelimbs (left) and hindlimbs (right) in *Pkd2l1::Cre; Rosa*Φ*DTR* mice 14 days after PBS (n=6) or DT (n=6) treatment.

D) Spontaneous rearing during a 90 min open field test in the center of the arena (left) or against the wall (right) in *Pkd2l1::Cre; Rosa*Φ*DTR* mice 14 days after PBS (n=8) or DT (n=6) treatment.

E) Speed during swimming task in *Pkd2l1::Cre; Rosa*Φ*DTR* mice 14 days after PBS (n=6) or DT (n=5) treatment.

F) Quantification of trunk angle between body axis and water line during swimming task in *Pkd2l1::Cre; Rosa*Φ*DTR* mice 14 days after PBS (n=6) or DT (n=5) treatment.

G) Representative images of Pkd2l1 staining in adult *Pkd2l1 +/+* and *Pkd2l1 -/-* animals.

H) Percentage of moving time in adult *Pkd2l1 +/+* (n=8) and *Pkd2l1 -/-* (n=9) during 90 min open field test.

I) Step cycle of forelimbs (top) and hindlimbs (bottom) in adult *Pkd2l1 +/+* (n=8) and *Pkd2l1 -/-* (n=10).

J) Quantification of foot placement errors (slips and falls) in the balance beam test with 2 cm (left) or 1 cm (right) beam width in adult *Pkd2l1 +/+* (n=8) and *Pkd2l1 -/-* (n=10).

K) Quantification of foot placement errors (slips and falls) in the horizontal ladder test with 1 cm (left) or 2 cm (right) rung distance in adult *Pkd2l1 +/+* (n=8) and *Pkd2l1 -/-* (n=10). Mean±SEM; paired t-test, ns p>0.05, * p<0.05, *** p < 0.001.

***Figure S4. Elimination of cilia from CSF-cN does not affect general locomotor activity and gait parameters***.

A and B) Representative electron microscopy images of CSF-cN (highlighted in yellow) in *control* (A) and Δ*cilia* (B) mice. Arrow point to the cilium.

C and D) Speed (C) and distance traveled (D) during a 90 min open field test in adult *control* (n=8) and Δ*cilia* (n=4) mice.

E) Speed during swimming task in adult *control* (n=10) and Δ*cilia* (n=9) mice.

F) Base of support of forelimbs (left) and hindlimbs (right) in adult *control* (n=10) and Δ*cilia* (n=9) mice.

G) Step length in adult *control* (n=10) and Δ*cilia* (n=9) mice (LF left forelimb, LH left hindlimb, RF right forelimb, RH right hindlimb).

Mean±SEM, paired t-test, ns p>0.05.

***Video S1. Representative runs of control, DT-treated***, Δ***cilium and Pkd2l1 -/- mice at the mouse walker***.

***Video S2. Representative runs of control, DT-treated***, Δ***cilium and Pkd2l1 -/- mice at the swim task***.

***Video S3. Representative runs of control, DT-treated***, Δ***cilium and Pkd2l1 -/- mice at the 1 cm diameter balance beam***.

***Video S4. Representative runs of control, DT-treated***, Δ***cilium and Pkd2l1 -/- mice at the 2 cm rung distance horizontal ladder***.

## References

Agduhr, E. (1922). Über ein zentrales Sinnesorgan (?) bei den Vertebraten. Z. Anat. Entwicklungsgesch. 66, 223–360.

Akay, T., and Murray, A.J. (2021). Relative Contribution of Proprioceptive and Vestibular Sensory Systems to Locomotion: Opportunities for Discovery in the Age of Molecular Science. Int. J. Mol. Sci. 22, 1–18.

Böhm, U.L., Prendergast, A., Djenoune, L., Figueiredo, S.N., Gomez, J., Stokes, C., Kaiser, S., Suster, M., Kawakami, K., Charpentier, M., et al. (2016). CSF-contacting neurons regulate locomotion by relaying mechanical stimuli to spinal circuits. Nat. Commun. 7, 1–8.

Buch, T., Heppner, F.L., Tertilt, C., Heinen, T.J. a J., Kremer, M., Wunderlich, F.T., Jung, S., and Waisman, A. (2005). A Cre-inducible diphtheria toxin receptor mediates cell lineage ablation after toxin administration. Nat. Methods 2, 419–426.

Daigle, T.L., Madisen, L., Hage, T.A., Valley, M.T., Knoblich, U., Larsen, R.S., Takeno, M.M., Huang, L., Gu, H., Larsen, R., et al. (2018). A Suite of Transgenic Driver and Reporter Mouse Lines with Enhanced Brain-Cell-Type Targeting and Functionality. Cell 174, 465–480.e22.

Dasen, J.S., and Jessell, T.M. (2009). Chapter Six Hox Networks and the Origins of Motor Neuron Diversity. In Current Topics in Developmental Biology, (Elsevier Inc.), pp. 169–200.

Dewitz, C., Pimpinella, S., Hackel, P., Akalin, A., Jessell, T.M., and Zampieri, N. (2018). Nuclear Organization in the Spinal Cord Depends on Motor Neuron Lamination Orchestrated by Catenin and Afadin Function. Cell Rep. 22, 1681–1694.

Djenoune, L., Khabou, H., Joubert, F., Quan, F.B., Nunes Figueiredo, S., Bodineau, L., Del Bene, F., Burcklé, C., Tostivint, H., and Wyart, C. (2014). Investigation of spinal cerebrospinal fluid-contacting neurons expressing PKD2L1: evidence for a conserved system from fish to primates. Front. Neuroanat. 8, 26.

Fidelin, K., Djenoune, L., Stokes, C., Prendergast, A., Gomez, J., Baradel, A., Del Bene, F., and Wyart, C. (2015). State-dependent modulation of locomotion by GABAergic spinal sensory neurons. Curr. Biol. 25, 3035–3047.

Grillner, S., and El Manira, A. (2020). Current principles of motor control, with special reference to vertebrate locomotion. Physiol. Rev. 100, 271–320.

Grillner, S., and Jessell, T.M. (2009). Measured motion: searching for simplicity in spinal locomotor networks. Curr. Opin. Neurobiol. 19, 572–586.

Gruner, J.A., and Altman, J. (1980). Swimming in the rat: Analysis of locomotor performance in comparison to stepping. Exp. Brain Res. 40, 374–382.

Haycraft, C.J., Zhang, Q., Song, B., Jackson, W.S., Detloff, P.J., Serra, R., and Yoder, B.K. (2007). Intraflagellar transport is essential for endochondral bone formation. Development 134, 307–316.

Horio, N., Yoshida, R., Yasumatsu, K., Yanagawa, Y., Ishimaru, Y., Matsunami, H., and Ninomiya, Y. (2011). Sour Taste Responses in Mice Lacking PKD Channels. PLoS One 6, e20007.

Hubbard, J.M., Böhm, U.L., Prendergast, A., Tseng, P.-E.B., Newman, M., Stokes, C., and Wyart, C. (2016). Intraspinal Sensory Neurons Provide Powerful Inhibition to Motor Circuits Ensuring Postural Control during Locomotion. Curr. Biol. 26, 2841–2853.

Jalalvand, E., Robertson, B., Tostivint, H., Wallén, P., and Grillner, S. (2016). The Spinal Cord Has an Intrinsic System for the Control of pH. Curr. Biol. 26, 1346–1351.

Jalalvand, E., Robertson, B., Wallén, P., and Grillner, S. (2016). Ciliated neurons lining the central canal sense both fluid movement and pH through ASIC3. Nat. Commun. 7, 10002.

Koch, S.C., Acton, D., and Goulding, M. (2018). Spinal Circuits for Touch, Pain, and Itch. Annu. Rev. Physiol. 80, 189–217.

Kolmer, W. (1921). Das „Sagittalorgan” der Wirbeltiere. Z. Anat. Entwicklungsgesch. 60, 652–717.

Lavin, T.K., Jin, L., Lea, N.E., and Wickersham, I.R. (2020). Monosynaptic Tracing Success Depends Critically on Helper Virus Concentrations. Front. Synaptic Neurosci. 12.

Li, Y., Stam, F.J., Aimone, J.B., Goulding, M., Callaway, E.M., and Gage, F.H. (2013). Molecular layer perforant path-associated cells contribute to feed-forward inhibition in the adult dentate gyrus. Proc. Natl. Acad. Sci. 110, 9106–9111.

Madisen, L., Mao, T., Koch, H., Zhuo, J.M., Berenyi, A., Fujisawa, S., Hsu, Y.W.A., Garcia, A.J., Gu, X., Zanella, S., et al. (2012). A toolbox of Cre-dependent optogenetic transgenic mice for light-induced activation and silencing. Nat. Neurosci. 15, 793–802.

Madisen, L., Zwingman, T.A., Sunkin, S.M., Oh, S.W., Zariwala, H.A., Gu, H., Ng, L.L., Palmiter, R.D., Hawrylycz, M.J., Jones, A.R., et al. (2010). A robust and high-throughput Cre Repooting and characterization. Nat Neurosci 13, 133–140.

Mathis, A., Mamidanna, P., Cury, K.M., Abe, T., Murthy, V.N., Mathis, M.W., and Bethge, M. (2018). DeepLabCut: markerless pose estimation of user-defined body parts with deep learning. Nat. Neurosci. 21, 1281–1289.

Mendes, C.S., Bartos, I., Márka, Z., Akay, T., Márka, S., and Mann, R.S. (2015). Quantification of gait parameters in freely walking rodents. BMC Biol. 13, 1–11.

Orts-Del’Immagine, A., Seddik, R., Tell, F., Airault, C., Er-Raoui, G., Najimi, M., Trouslard, J., and Wanaverbecq, N. (2016). A single polycystic kidney disease 2-like 1 channel opening acts as a spike generator in cerebrospinal fluid-contacting neurons of adult mouse brainstem. Neuropharmacology 101, 549–565.

Orts-Del’Immagine, A., and Wyart, C. (2017). Cerebrospinal-fluid-contacting neurons. Curr. Biol. 27, R1198–R1200.

Orts-Del’Immagine, A., Cantaut-Belarif, Y., Thouvenin, O., Roussel, J., Baskaran, A., Langui, D., Koëth, F., Bivas, P., Lejeune, F.-X.X., Bardet, P.-L.L., et al. (2020). Sensory Neurons Contacting the Cerebrospinal Fluid Require the Reissner Fiber to Detect Spinal Curvature In Vivo. Curr. Biol. 30, 827–839.e4.

Pazour, G.J., Dickert, B.L., Vucica, Y., Seeley, E.S., Rosenbaum, J.L., Witman, G.B., and Cole, D.G. (2000). Chlamydomonas IFT 88 and Its Mouse Homologue, Polycystic Kidney Disease Gene Tg 737, Are Required for Assembly of Cilia and Flagella. J. Cell Biol. 151, 709–718.

Petreanu, L., Huber, D., Sobczyk, A., and Svoboda, K. (2007). Channelrhodopsin-2–assisted circuit mapping of long-range callosal projections. Nat. Neurosci. 10, 663–668.

Pocratsky, A.M., Shepard, C.T., Morehouse, J.R., Burke, D.A., Riegler, A.S., Hardin, J.T., Beare, J.E., Hainline, C., States, G.J., Brown, B.L., et al. (2020). Long ascending propriospinal neurons provide flexible, context-specific control of interlimb coordination. Elife 9, 1–24.

Rossignol, S., Dubuc, R., and Gossard, J.-P. (2006). Dynamic Sensorimotor Interactions in Locomotion. Physiol. Rev. 86, 89–154.

Rudomin, P., and Schmidt, R.F. (1999). Presynaptic inhibition in the vertebrate spinal cord revisited. Exp. Brain Res. 129, 1–37.

Stepien, A.E., Tripodi, M., and Arber, S. (2010). Monosynaptic rabies virus reveals premotor network organization and synaptic specificity of cholinergic partition cells. Neuron 68, 456–472.

Sternberg, J.R., Prendergast, A.E., Brosse, L., Cantaut-Belarif, Y., Thouvenin, O., Orts-Del’Immagine, A., Castillo, L., Djenoune, L., Kurisu, S., McDearmid, J.R., et al. (2018). Pkd2l1 is required for mechanoception in cerebrospinal fluid-contacting neurons and maintenance of spine curvature. Nat. Commun. 9, 1–10.

Stoeckel, M.-E., Uhl-Bronner, S., Hugel, S., Veinante, P., Klein, M.-J., Mutterer, J., Freund-Mercier, M.-J., and Schlichter, R. (2003). Cerebrospinal fluid-contacting neurons in the rat spinal cord, a γ-aminobutyric acidergic system expressing the P2X2 subunit of purinergic receptors, PSA-NCAM, and GAP-43 immunoreactivities: Light and electron microscopic study. J. Comp. Neurol. 457, 159–174.

Tuthill, J.C., and Azim, E. (2018). Proprioception. Curr. Biol. 28, R194–R203.

U, W. (1996). On the role of recurrent inhibitory feedback in motor control. Prog. Neurobiol. 49, 517–587.

Vígh, B., Manzanoe Silva, M.J., Frank, C.L., Vincze, C., Czirok, S.J., Szabó, a., Lukáts, a., and Szél, a. (2004). The system of cerebrospinal fluid-contacting neurons. Its supposed role in the nonsynaptic signal transmission of the brain. Histol. Histopathol. 19, 607–628.

Wickersham, I.R., Lyon, D.C., Barnard, R.J.O.O., Mori, T., Finke, S., Conzelmann, K.-K.K., Young, J.A.T.T., and Callaway, E.M. (2007). Monosynaptic restriction of transsynaptic tracing from single, genetically targeted neurons. Neuron 53, 639–647.

Witts, E.C., Zagoraiou, L., and Miles, G.B. (2014). Anatomy and function of cholinergic C bouton inputs to motor neurons. J. Anat. 224, 52–60.

Wu, M.-Y., Carbo-Tano, M., Mirat, O., Lejeune, F.-X., Roussel, J., Quan, F.B., Fidelin, K., and Wyart, C. (2021). Spinal sensory neurons project onto the hindbrain to stabilize posture and enhance locomotor speed. Curr. Biol. 31, 3315–3329.e5.

Ye, W., Chang, R.B., Bushman, J.D., Tu, Y.-H., Mulhall, E.M., Wilson, C.E., Cooper, A.J., Chick, W.S., Hill-Eubanks, D.C., Nelson, M.T., et al. (2015). The K + channel K IR 2.1 functions in tandem with proton influx to mediate sour taste transduction. Proc. Natl. Acad. Sci. 201514282.

Zagoraiou, L., Akay, T., Martin, J.F., Brownstone, R.M., Jessell, T.M., Miles, G.B., Thomas, M., and Miles, G.B. (2009). A Cluster of Cholinergic Premotor Interneurons Modulates Mouse Locomotor Activity. Neuron 64, 645–662.

Zampieri, N., Jessell, T.M., and Murray, A.J. (2014). Mapping Sensory Circuits by Anterograde Transsynaptic Transfer of Recombinant Rabies Virus. Neuron 81, 766–778.

