## Supplemental information for "The role of intraspinal sensory neurons in the control of quadrupedal locomotion"

**Supplemental Information Titles and Legend**

***Figure S1. Specificity of CSF-cN genetic targeting.***

A) Representative image of Pkd2l1$^{+}$; nuclear GFP$^{+}$ neurons around the central canal of *Pkd2l1::Cre; RosaΦHTB* mice.

B) Quantification of Pkd2l1$^{+}$; nuclear GFP$^{+}$ neurons at cervical, thoracic, and lumbar spinal segments (cervical 182/220 cells, thoracic 198/230 cells, lumbar 179/209 cells, n=4; Mean±SEM).

C) Tomato expression in the brain of P7 *Pkd2l1::Cre; RosaΦtdTomato* mice (cortex, Ctx; ventricular zone, VZ; cerebellum, CB). Scale bar: 1mm.

D) Representative pictures of synaptophysin-tdTomato synapses on ChAT$^{+}$ V0c, MMC, and LMC neurons at cervical and lumbar segments of P7 *Pkd2l1::Cre; Ai34* mice.

***Figure S2. Rabies monosynaptic tracing from CSF-cN.***

A) Digital reconstruction of BFP$^{+}$; RV$^{+}$ starter cells (blue) position. Each plot represents one animal.

B) Digital reconstruction of second order cells (red) position. Each plot represents one animal.

C) Cross correlation analysis of starter cells (left) and second order cells (right) positional coordinates. The scale indicates correlation values.

D) Number of starter cells (blue), second order cells (red) and second order CSF-cN in individual rabies tracing experiments (right, black).

E) Proportion of VGLUT2$^{+}$ and VGAT$^{+}$ second order neurons (excluding CSF-cN).

Mean ± SEM, paired t-test, * $p<0.05$.

***Figure S3. Behavioral analysis of CSF-cN ablated and Pkd2l1 mutant mice.***

A) Representative images of Pkd2l1 staining in adult *Pkd2l1::Cre; RosaΦDTR* mice 7 days after PBS (left) or DT-injection (right).

B) Percentage of Pkd2l1$^{+}$ cells in respect to the PBS control at 7 days, 11 days and 60 days after DT treatment in adult *Pkd2l1::Cre; RosaΦDTR* mice.

C) Base of support of forelimbs (left) and hindlimbs (right) in *Pkd2l1::Cre; RosaΦDTR* mice 14 days after PBS (n=6) or DT (n=6) treatment.

***Video S2. Representative runs of control, DT-treated, Δcilium and Pkd2l1 -/- mice at the swim task.***

***Video S3. Representative runs of control, DT-treated, Δcilium and Pkd2l1 -/- mice at the 1 cm diameter balance beam.***

***Video S4. Representative runs of control, DT-treated, Δcilium and Pkd2l1 -/- mice at the 2 cm rung distance horizontal ladder.***

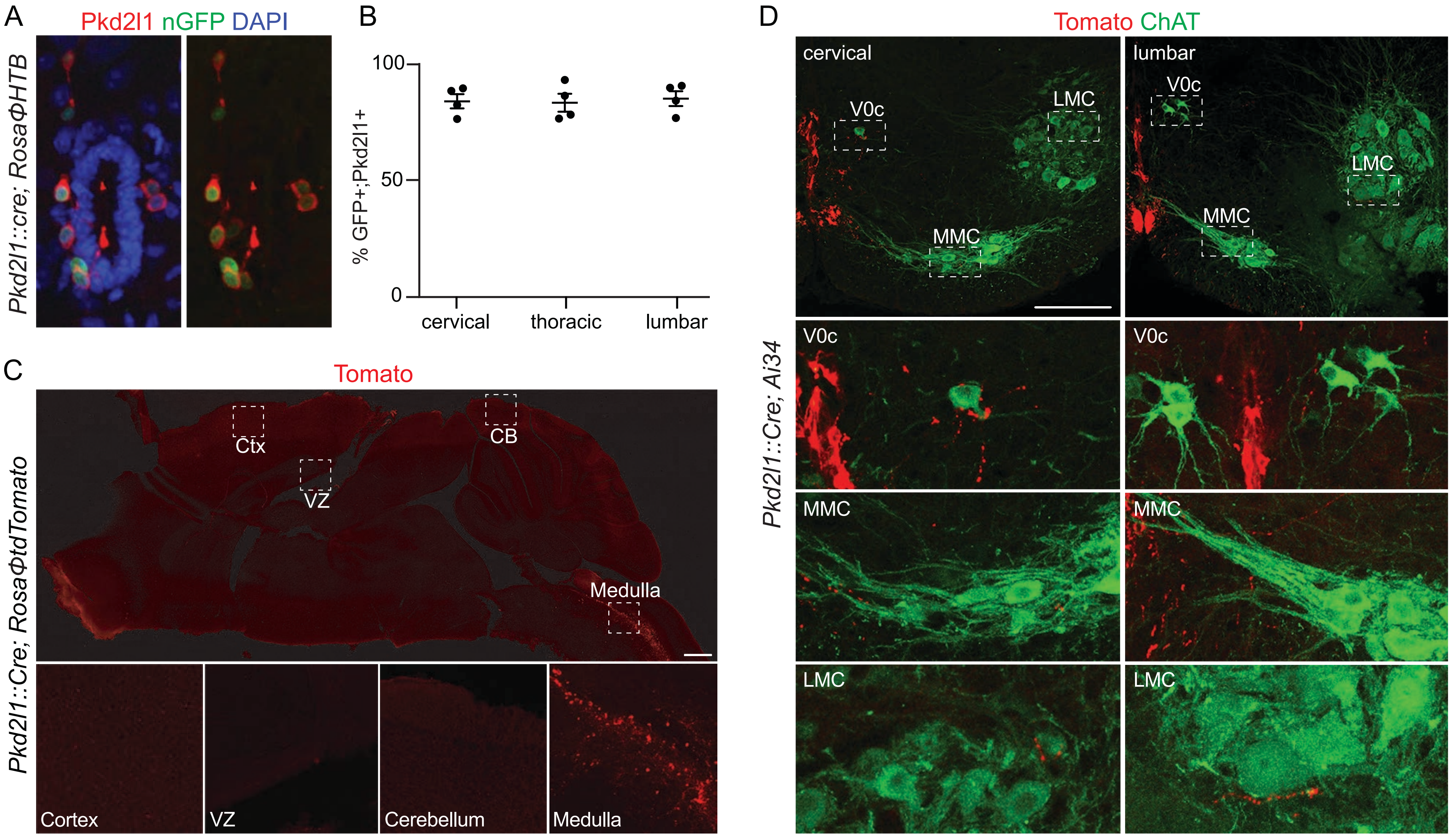

A
Pkd2l1 nGFP DAPI
Pkd2l1::cre; RosaΦHTB
B
100
50
0
% GFP+;Pkd2l1+
cervical
thoracic
lumbar
C
Tomato
Pkd2l1::Cre; RosaΦtdTomato
Ctx
CB
VZ
Medulla
Cortex
VZ
Cerebellum
Medulla
D
Tomato ChAT
Pkd2l1::Cre; Ai34
cervical
lumbar
V0c
LMC
MMC
V0c
MMC
LMC
V0c
V0c
MMC
MMC
LMC
LMC


Figure S1

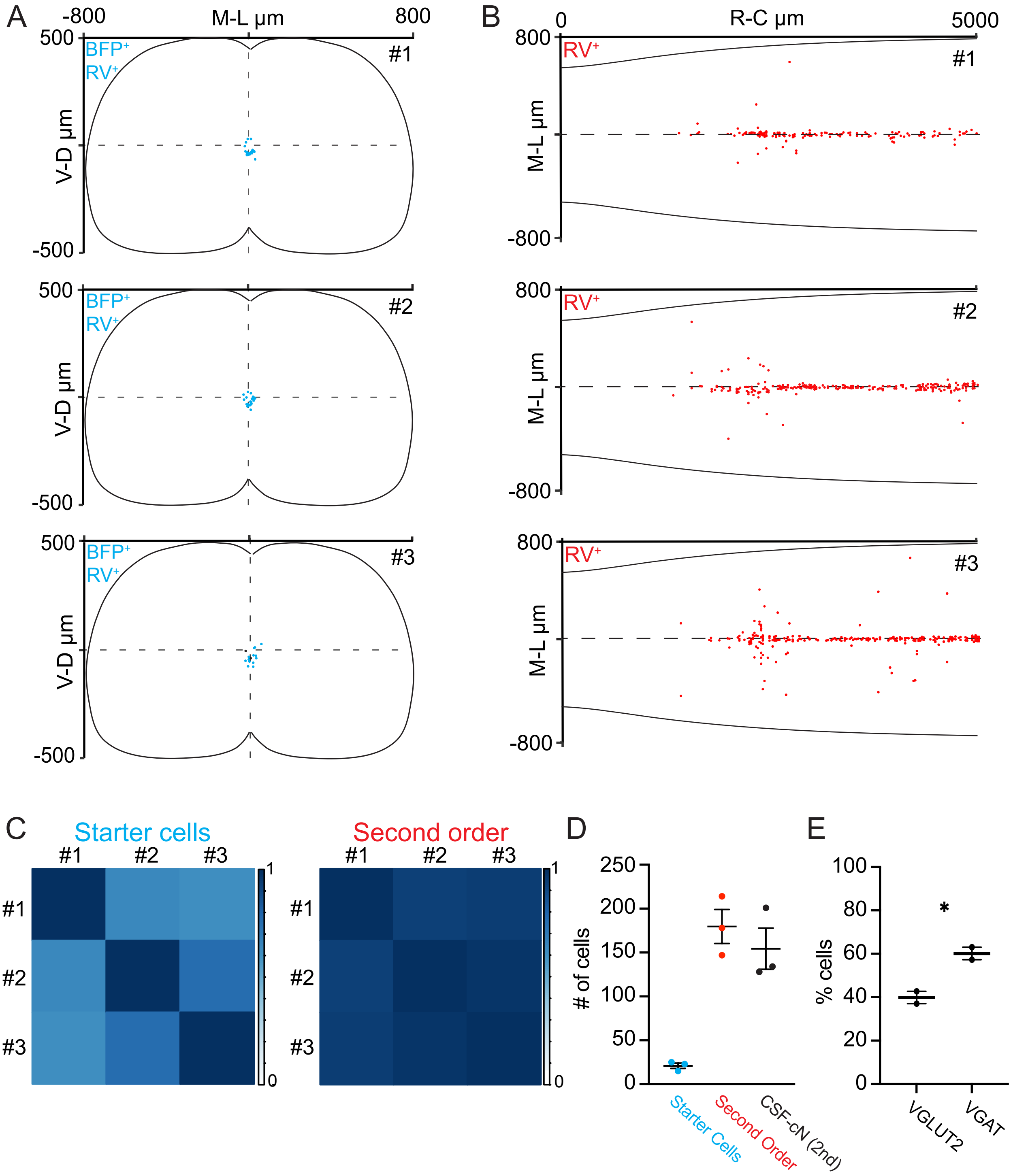
A
-800
M-L µm
800
500
-500
V-D µm
BFP+
RV+
#1
#2
#3
B
0
R-C µm
5000
800
-800
M-L µm
RV+
C
Starter cells
Second order
#1
#2
#3
1
0
D
### of cells
250
200
150
100
50
0
Starter Cells
Second Order
CSF-cN (2nd)
E
% cells
100
80
60
40
20
0
*
VGLUT2
VGAT

Figure S2

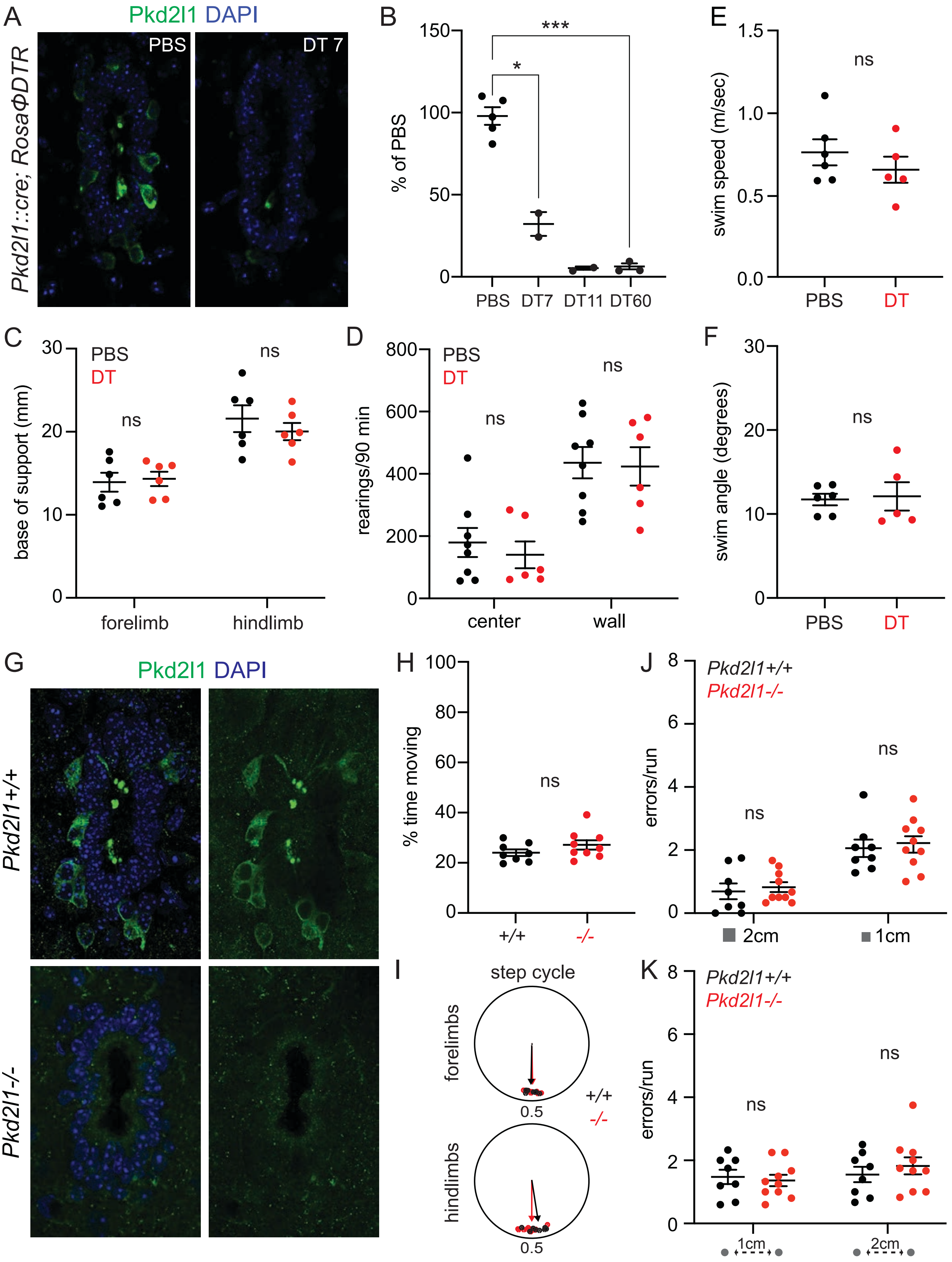
A
Pkd2l1 DAPI
PBS
DT 7
Pkd2l1::cre; RosaΦDTR
B
% of PBS
***
*
PBS DT7 DT11 DT60
E
swim speed (m/sec)
ns
PBS DT
C
base of support (mm)
PBS
DT
ns
forelimb hindlimb
D
rearings/90 min
center wall
F
swim angle (degrees)
G
Pkd2l1 DAPI
Pkd2l1+/+
Pkd2l1-/-
H
% time moving
+/+ -/-
J
errors/run
2cm
1cm
I
step cycle
forelimbs
hindlimbs
0.5
K
1cm
2cm

Figure S3

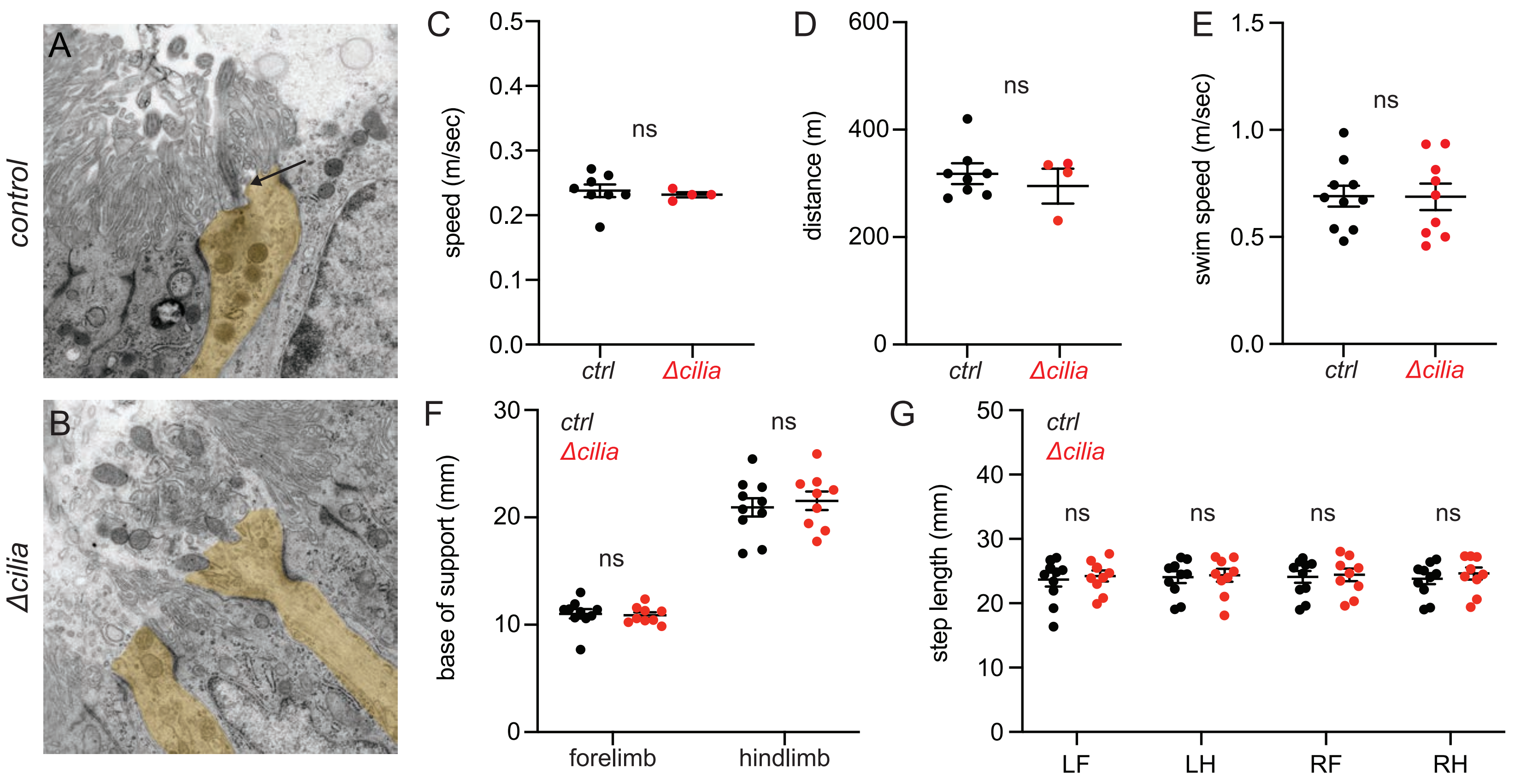
A
control
B
Δcilia
C
speed (m/sec)
0.0
0.1
0.2
0.3
0.4
0.5
ns
ctrl
Δcilia
D
distance (m)
0
200
400
600
ns
ctrl
Δcilia
E
swim speed (m/sec)
0.0
0.5
1.0
1.5
ns
ctrl
Δcilia
F
base of support (mm)
0
10
20
30
ctrl
Δcilia
ns
ns
forelimb
hindlimb
G
step length (mm)
0
10
20
30
40
50
ctrl
Δcilia
ns
ns
ns
ns
LF
LH
RF
RH

Figure S4
